# Drilling down hotspots of intraspecific diversity to bring them into on-ground conservation of threatened species

**DOI:** 10.1101/419788

**Authors:** Mauro Zampiglia, Roberta Bisconti, Luigi Maiorano, Gaetano Aloise, Antonino Siclari, Francesco Pellegrino, Giuseppe Martino, Alice Pezzarossa, Andrea Chiocchio, Chiara Martino, Giuseppe Nascetti, Daniele Canestrelli

## Abstract

Unprecedented rates of biodiversity loss rise the urgency for preserving species ability to cope with ongoing global changes. An approach in this direction is to target intra-specific hotspots of genetic diversity as conservation priorities. However, these hotspots are often identified by sampling at a spatial resolution too coarse to be useful in practical management of threatened species, hindering the long-appealed dialog between conservation stakeholders and conservation genetic researchers. Here, we investigated the spatial and temporal variation in species presence, genetic diversity, as well as potential risk factors, within a previously identified hotspot of genetic diversity for the endangered Apennine yellow bellied toad *Bombina pachypus*. Our results show that this hotspot is neither a geographically homogeneous nor a temporally stable unit. Over a time-window spanning 10-40 years since previous assessments, *B. pachypus* populations declined in large portions of its hotspot, and their genetic diversity levels decreased. Considering the demographic trend, genetic and epidemiological data, and models of current and future climatic suitability, populations at the extreme south of the hotspot area still qualify for urgent in-situ conservation actions, whereas northern populations would be better managed through a mix of in-situ and ex-situ actions. Our results emphasize that identifying hotspot of genetic diversity, albeit essential step, does not suffice to warrant on-ground conservation of threatened species. Hotspots should be analysed at finer geographic and temporal scales, to provide conservation stakeholders with key knowledge to best define conservation priorities, and to optimize resource allocation to alternative management practices.

## 1. Introduction

Unprecedented rates of biodiversity loss rise the urgency for preserving species ability to cope with the ongoing global changes (Crandall et al., 2000; Eizaguirre and Baltazar-Soares, 2014 and references therein; Hendry et al., 2010; Moritz, 2002). However, conservation policies have traditionally been oriented towards the protection of species diversity and habitats, rarely considering the evolutionary potential of individual populations within species (Laikre, 2010; Mace and Purvis, 2008; Mimura et al., 2017). A direct approach in this direction, which has been largely debated for decades, is to directly target populations with high levels of genetic diversity (Frankham, 2010). In fact, owing to the intimate links between genetic diversity and effective population size, these populations are, at the same time, less likely to be affected by the detrimental consequences of inbreeding depression and genetic drift, and more likely to warrant evolvability and adaptive potential (Allendorf et al., 2012; Frankham, 2010; Lanfear, et al., 2014). When different populations, all with relatively high levels of genetic diversity, are concentrated in the same area it is possible to define an intra-specific hotspot of genetic diversity (Hampe and Petit, 2005; Hewitt, 2000, 2004; Petit et al., 2003), clearly representing a conservation priority. Importantly, as pointed out by Pérez-Espona and colleagues (2017), genetic diversity data useful to detect hotspots, are now available for a large amount of taxa, at least within some intensively studied areas. However, in spite of both their extreme value and data availability, hotspots of genetic diversity have been largely ignored in the context of biodiversity management strategies, highlighting an existing gap between conservation geneticists’ achievements and conservation stakeholders’ priorities (Pérez-Espona and ConGRESS Consortium, 2017; Vernesi et al., 2008).

Traditionally, hotspots have been identified over large areas (e.g., considering the entire range of a species) using the analytical tools and the sampling scheme of phylogeography (Hewitt, 2011), providing therefore a spatial resolution that can be hardly considered in practical conservation exercises (Cañadas and Vázquez, 2014). Consequently, important questions for on-ground management practices, still remain largely unexplored. Can hotspot be considered as homogeneous regional units for management purposes? Are all populations ‘created’ equal within hotspot areas? Or, in order to achieve optimal resource allocation and maximum chance of success, should populations be prioritized as well? Answering these open questions has major practical implications for the apportionment of financial budgets among alternative management practices, as well as for the design of protected areas.

Here, we demonstrate the importance of addressing these questions, in a collaborative framework between academic researchers and conservation authorities (governmental, protected area managers). We investigated spatial and temporal patterns of variation in species presence, genetic diversity, as well as potential risk factors, within a previously identified hotspot of genetic diversity for the endangered Apennine yellow bellied toad *Bombina pachypus*. Indeed, substantial previous knowledge available for this species makes it especially suitable to reach the study aim. The Apennine yellow-bellied toad is an amphibian endemic to the Italian peninsula, whose hotspot of genetic diversity has been identified at the southernmost portion of its range (Canestrelli et al., 2006). Extensive population declines in the last decades have raised severe concern for the conservation status of this species (Barbieri et al., 2004), that is now listed as endangered in the Red List of IUCN (Andreone et al., 2009). Previous studies have suggested the potential conservation value of the hotspot area for this species, showing that: i) *B. pachypus* populations are facing a dramatic demographic decline in the whole range, except for this area (Barbieri et al., 2004); ii) genetic diversity levels in this area are by far higher than in the rest of the range (Canestrelli et al., 2006); and iii) populations from this area are reported as viable despite the long-term co-occurrence of the ‘killer fungus’ *Batrachochytrium dendrobatidis* (Canestrelli et al., 2013), considered one of the main drivers of the decline (Berger et al., 1998; Stagni et al., 2004).

To achieve our goals, we integrated previous knowledge with the results of three experimental steps. First, we surveyed the whole region to assess whether occurrence pattern of populations has remained stable over time, in the whole area or in parts of it. Second, we assessed the genetic structure of populations, quantified their levels of genetic diversity, and compared them with those of populations sampled in the same area in the past. Third, we performed an analysis of current potential risk factors by focusing on the present-day occurrence of the chytrid pathogen *B. dendrobatidis*, as well as by estimating variation in bioclimatic suitability of the hotspot area, from current to future climatic scenarios.

## 2. Material and Methods

### 2.1. Species presence data

Barbieri and colleagues (2004) indicated southern Italy (Calabria administrative region) as the only area within the distribution range where the yellow bellied toad was not facing demographic declines at the end of 1990s. In order to confirm this finding and/or to characterize spatial and temporal patterns of variation, we carried out extensive field surveys within this region. Field activities were carried out from April to October, corresponding with the reproductive season of the species, from 2013 to 2016. Field activities and sampling procedures were approved by the Italian Ministry of the Environment (protocol numbers: 0042634/PNM, and 0007727/PNM).

We screened peer-reviewed and grey literature, and museum collections for records of presence of the toad in Calabria. We retained only 56 records with detailed information about location and year of observation and we divided the region into 4 areas where most of the records are concentrated: Catena Costiera, Sila, Serre and Aspromonte (Figure 1). The boundaries of these 4 areas were defined arbitrarily, although roughly based on mountain ranges.

**Figure 1.**
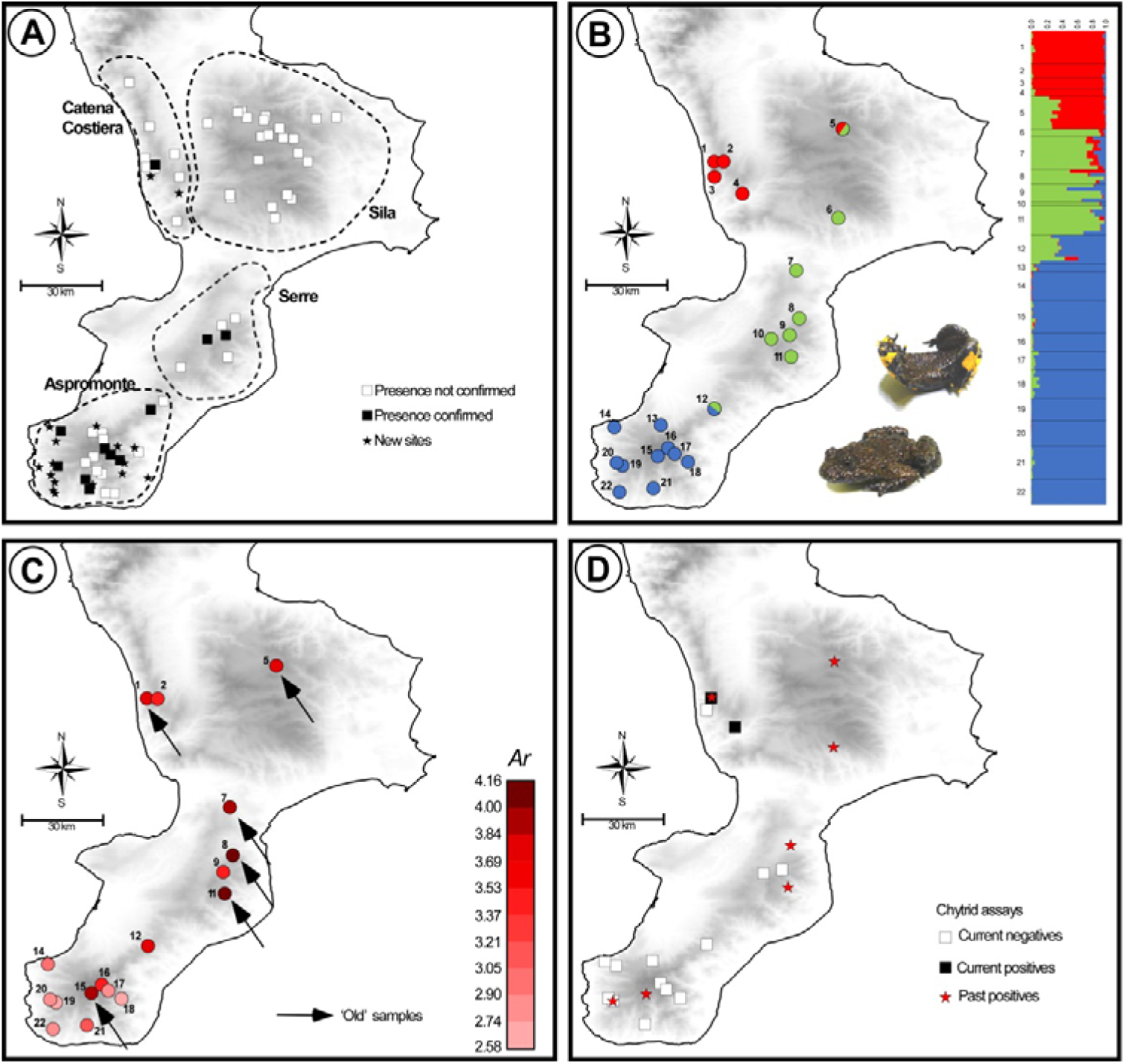
Spatial and temporal patterns of variation in species presence, genetic diversity, and chytrid occurrence, for the Apennine yellow-bellied toad in southern Italy (Calabria region). Sources of historical data and samples are reported within the main text. A: Sites of past and current presence of *B. pachypus* in the area; dashed shapes show the four subregions considered in this study. B: Results of the Bayesian clustering analysis used to assess the population genetic structure of *B. pachypus* in the area; admixture proportions of each sampled individual for the 3 genetic clusters identified as the best clustering option by the method implemented in TESS. C: Geographic variation in estimates of genetic diversity (A_r_: allelic richness), among populations of *B. pachypus* sampled either for the present study or in the past. D: Past and current presence of the chytrid pathogen *B. dendrobatidis* among the sampled populations of *B. pachypus* from the study area.

Each site of historical presence was visited three times during the whole campaign. We considered the species as currently present in a site when at least one evidence of its presence (either adults, or tadpoles, or clutches) was observed in at least one visit. Field research was also opportunistically extended to other 80 non-reported but potentially suitable sites in the surroundings, in order to search for new records.

### 2.2 Genetic diversity and population structure

Population genetic structure and diversity were assessed by genotyping 130 individuals from 22 sites, ranging over the whole region (Table 1 and Figure 1). Individuals from 15 of these sites (“new” samples in Table 1) were sampled by toe-clipping after anaesthetization in a 0.1% solution of MS222 (3-aminobenzoic acid ethyl ester). All individuals were immediately released at the collection sites, while tissue samples were stored in 95% alcohol. Tissue samples from the remaining 7 sites (“old” samples in Table 1) were collected during previous studies carried out from 1978 to 2006 (see Canestrelli et al., 2006, 2013 and references therein), and were stored in 95% alcohol.

**Table 1.**
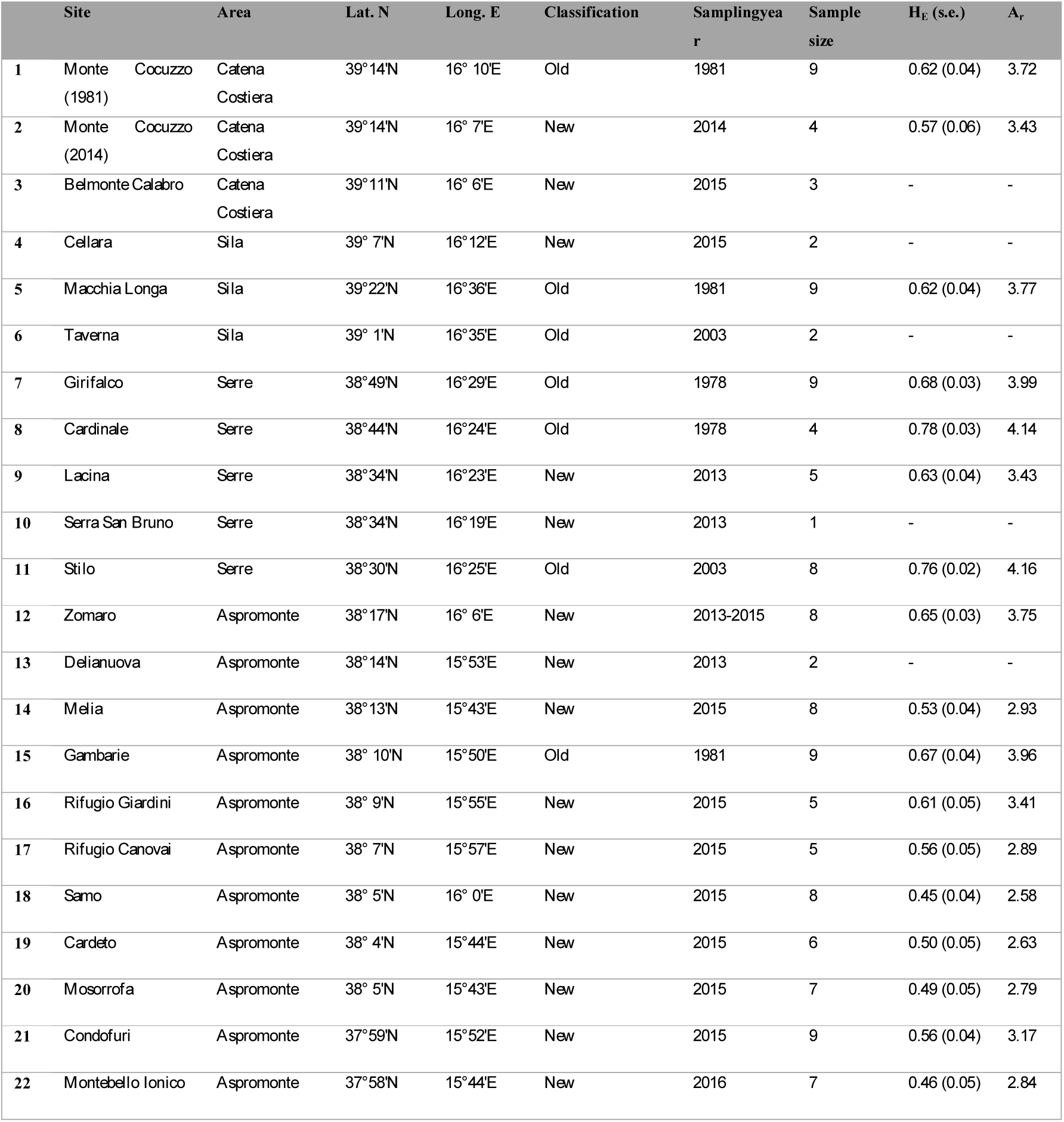
Geographic location, sample size, sampling year, and estimates of genetic diversity, for the 22 populations of *B. pachypus* sampled for the analysis of genetic variation in southern Italy. Sampling areas are defined according to Figure 1. H_E_: unbiased expected heterozygosity; A_r_: allelic richness.

Genomic DNA was extracted using the ZR Genomic DNA™ -Tissue MiniPrep kit (Zymoresearch) following manufacturer’s instructions. We initially tested 11 microsatellite loci that previously proved to amplify in the sister species *B. variegata*: Bv11.7, Bv32.7, B13, B14, F22, 1A, 5F, 8A, 9H, 10F, 12F (Hauswaldt et al., 2007; Nürnberger et al., 2003; Stuckas and Tiedemann 2006). We excluded from the analyses the loci Bv32.7 (failed the amplification step for our samples), B13 (yielded inconsistent reactions), B14 (yielding missing data in 26% of the individuals), and 12F (monomorphic in preliminary trials based on 30 randomly selected individuals). PCR conditions for the 7 loci analysed in this study, along with fluorescent dyes used to label forward primers and multiplex assembly are shown in Supplementary material. Fragment analysis of PCR products was performed by Macrogen Inc. on an ABI 3730xl Genetic Analyser (Applied Biosystems) with a 400HD size standard.

Allele calling was performed with GENEMAPPER^®^ 4.1 checking electropherogram by eye. Microsatellite dataset was then analysed with MICRO-CHECKER 2.2.3(Van Oosterhout et al., 2004) to test for the presence of null alleles, large alleles drop-out or scoring errors. We used GENEPOP 4.5.1 (Raymond and Rousset, 1995; Rousset, 2008) to test for departures from Hardy-Weinberg equilibrium and for genotypic linkage disequilibrium. Population genetic diversity was estimated as expected heterozigosity (H_e_) and allelic richness (A_r_) using GENETIX4.05.2 (Belkhir et al., 1996) and FSTAT 2.9.3.2 (Goudet, 1995, 2001), respectively. These analyses were carried out excluding sites where we sampled less than four individuals.

Genetic population structure was inferred by means of the Bayesian clustering algorithm implemented in TESS 2.3.1 (Chen et al., 2007),with the admixture model, under the conditional autoregressive model (CAR), and the geographical coordinates of individuals as prior information (Durand et al., 2009).TESS was used as clustering method since it has been shown to perform better than similar methods when the number loci is limited and/or when genetic structure is shallow (Chen et al., 2007; François and Durand, 2010). We performed a set of preliminary analyses with 20000 runs, discarding the first 5000 as burn-in, and with 10 replicates for each value of K from 2 to 10 to test model performance. For the final analysis we ran 100 replicates for each value of K from 2 to 6, with 100000 steps and discarding the first 50000 as burn-in. The spatial interaction parameter was set to the default value (0.6) and the option of update this parameter was activated. The clustering model that best fitted the data was inferred by the deviance information criterion (DIC), averaging the DIC values over the 100 replicates for each K and selecting the K value at which the average DIC reached a plateau. The 10 runs with the lowest DIC values for the inferred K were finally selected, and the estimated admixture proportions were averaged over them using CLUMPP 1.1.2 (Jakobsson and Rosenberg, 2007).

### 2.3 Pathogen assay

Skin swabs were collected from 105 individuals collected in 15 sites (Figure 1). Genomic DNA was extracted from swabs following the protocol of Boyle and colleagues (2004)as modified by Zampiglia and colleagues (2013). The molecular diagnostic assay was conducted in 25 µl reaction volumes using a nested PCR protocol as developed by Goka and colleagues (2009). The assay was performed once for each sample and it included a positive (DNA extracted from *B. dendrobatidis* zoospores JEL423, kindly provided by Prof. Joyce Longcore) and a negative (DNA-free distilled water) control. PCR products were loaded on a 1% agarose gel and checked for the diagnostic band at approximately 300 bp size to be considered as *B. dendrobatidis* positive. A geographic representative subset of samples (20%) was analysed in duplicate.

### 2.4 Changes in bioclimatic suitability

We calibrated species distribution models (SDM) with a dataset of 361 occurrences covering the entire distribution range of the species and a set of bioclimatic variables. The occurrences were obtained by pooling the data collected during the field campaign (this study) and different data sources: Stoch (2000-2005), Global Biodiversity Information Facility (www.GBIF.org), Observado (www.observation.org) and Canestrelli and colleagues (2006). The dataset was filtered to remove duplicated records and data georeferenced with uncertainty. To limit spatial autocorrelation, we thinned the raw occurrences using the SPTHIN R package (Aiello-Lammens et al., 2015), obtaining five alternative calibration subsets with 182 occurrence data (see Supplementary material for details). Considering the same data sources available for the presence, we collected all data available for the same study area for species of reptiles and amphibians. We collected a total of 7614 records that we used in model calibration as background points in order to limit the effects of the existing sampling bias (Ranc et al., 2017).

The bioclimatic variables for current climate were obtained from Hijmans and colleagues (2005). Following a variance inflation factor (VIF) analysis on the original 18 bioclimatic variables, we considered in the analyses only the following 7 variables with VIF < 5: annual mean temperature, mean diurnal range, temperature seasonality, mean temperature of wettest quarter, mean temperature of driest quarter, precipitation of wettest month, and precipitation seasonality. For the future climate, we considered the same 7 variables calculated under three general circulation models (CCSM4, MIROC-ESM, MPI-ESM-P) and two emission scenarios (RCP2.6 and RCP8.5) adopted by the IPCC5 (Intergovernmental Panel on Climate Change).

Using an ensemble forecasting approach (Araujo and New, 2007), we calibrated the SDM considering the current climate, the five thinned occurrence sets, the set of background points, and the following five algorithms as implemented in the BIOMOD2 R packages (Thuiller et al., 2009): generalized linear model (GLM); generalized additive model (GAM); generalized boosted models (GBM); multivariate adaptive regression spline (MARS), and maximum entropy (MAXENT).

For model evaluation, we considered again five sets of points, each including the 179 occurrence points excluded through the thinning procedure from the calibration datasets and 7488 random background points (maintaining the same prevalence as in the calibration dataset). For each model, we measured the area under the curve (AUC) of the receiver operating characteristic (ROC) curve. Models with AUC value greater than 0.7 (Swets, 1988) were projected under the current and future climate over the entire peninsular Italy. Then, for each projection we obtained a final consensus SDM, calculated as the weighted average of all available models (weights for each model based on the respective AUC score, as in Marmion et al.,2009).

## 3. Results

### 3.1 Species presence in space and time

We found evidence of presence of *B. pachypus* in 28 localities out of the original 136 (21%), including 11 sites of historical presence (out of the original 56) and 17 new records. The highest number of historical sites with confirmed presence was found in the Aspromonte (8 out of 19; 42%), followed by the Serre (2 out of 6; 33%) and the Catena Costiera (1 out of 8; 13%). No presence was confirmed for the Sila (0 presences out of the original 23). The 17 new records of presences were unevenly distributed among regions, with 15 new sites found in the Aspromonte, 2 sites found in the Catena Costiera, and no new site found in Sila and Serre regions. The Aspromonte region was also the area with the highest number of individual observed per site/day, with an average of 21.6 individuals (s.d. 18.3) compared to 3.5 individuals (s.d. 1.9) observed on average in the other regions.

### 3.2 Genetic diversity and population structure

The number of alleles observed at each microsatellite locus was as follows: 1A = 11, 8A = 13, 5F = 12, 10F = 12, 9H = 25, Bv11 = 3 and F22 = 9. All loci were used for downstream analyses, since none of them showed evidence of null alleles in more than one sampling site. Missing data accounted to 3.7% over the whole dataset. No evidence of genotypic linkage disequilibrium between loci was observed, and no departures from the Hardy-Weinberg equilibrium was detected after the Bonferroni correction was applied (Rice, 1989).

Expected heterozigosity ranged from 0.45 (site 18) to 0.78 (site 8), while allelic richness ranged from 2.58 (site 18) to 4.16 (site 11; Figure 1C). Estimates of both the expected heterozigosity and the allelic richness for each sampled site are shown in Table 1. Among the ‘old’ samples, the highest values of both heterozigosity and allelic richness were observed within Serre region, whereas among the ‘new’ samples, samples with the highest values of genetic diversity were observed within Aspromonte region. Although the location and size of our samples prevented us from carrying out quantitative temporal comparisons, it is worth noting that ‘old’ sample systematically yielded higher values of genetic diversity when compared with the nearest ‘new’ samples from the same area (see Table 1).

Bayesian clustering analysis of population structure revealed the presence of three main population clusters within the study area. Pie-charts displaying the contribution of the three clusters to the genetic pool of each site and bar-plots indicating individual admixture proportions are shown in Figure 1B. The three clusters showed a clear geographic structure along the north-south axis: one was restricted to the north, in Catena Costiera and northern Sila, one had a central distribution, ranging from southern Silato Serre, and the third one was restricted to the south, in Aspromonte. Evidence of admixture between clusters was observed in some individuals, particularly from sites located in intermediate areas (sites 5 and12).

### 3.3 Occurrence of *Batrachochytrium dendrobatidis*

The diagnostic tests for *B. dendrobatidis* occurrence yielded positive results for the presence of the pathogen on the skin of 3 individuals from 2 sites in the Catena Costiera. Instead, and in contrast to previous assessments (Canestrelli et al.,2013), no positive tests were observed for individuals sampled in the Serre and Aspromonte (Figure 1D). All the analyses carried out in duplicate yielded fully consistent results.

### 3.4 Species distribution modelling

We calibrated 25 models (5 presence-background sets for 5 modelling techniques) with, on average, good model performance (mean AUC = 0.80; s.d. AUC = 0.04). GBM was the model with the highest AUC values (mean AUC = 0.86) and the GLM the one with the lowest (mean AUC = 0.76). Temperature was by far the most important bioclimatic factor for explaining the distribution of the yellow-bellied toad, with mean temperature of the driest quarter, mean temperature of the wettest quarter, and annual mean temperature being the most important variables, followed by the two precipitation variables (Table 2).

**Table 2.** Variable importance for the different algorithms. Values in the table are the average across all modeling runs. A value of 0 assumes no influence of the variable on the model; the higher the value, the more influence the variable has on the model (maximum = 1; Thuiller, Lafourcade, Engler and Araújo, 2009). Climatic variables defined in Hijmans and colleagues 2005.

|  | GLM | GBM | GAM | MARS | MAXENT |
| --- | --- | --- | --- | --- | --- |
| **Annual Mean Temperature** | 0.2776 | 0.0398 | 0.638 | 0.3724 | 0.0526 |
| **Mean Diurnal Range** | 0.0564 | 0.1744 | 0.046 | 0.0518 | 0.0574 |
| **Temperature Seasonality** | 0.0892 | 0.1094 | 0.235 | 0.1472 | 0.1038 |
| **Mean Temperature of Wettest Quarter** | 0.2074 | 0.2478 | 0.2064 | 0.1662 | 0.2264 |
| **Mean Temperature of the Driest Quarter** | 0.1822 | 0.2006 | 0.6012 | 0.3826 | 0.2162 |
| **Precipitation of the Wettest Month** | 0.1612 | 0.0892 | 0.145 | 0.137 | 0.1118 |
| **Precipitation Seasonality** | 0.0966 | 0.037 | 0.213 | 0.1814 | 0.1444 |

Regardless of the climate scenario considered, the future climate suitability is predicted to decrease substantially (Figure 2), especially in the north and central Apennines, where bioclimatic suitability drops dramatically in some areas. A much higher climatic stability in time is predicted for the southern part of the distribution range, where however the models predict a future range shift towards higher elevations (Figure 2).

**Figure 2.**
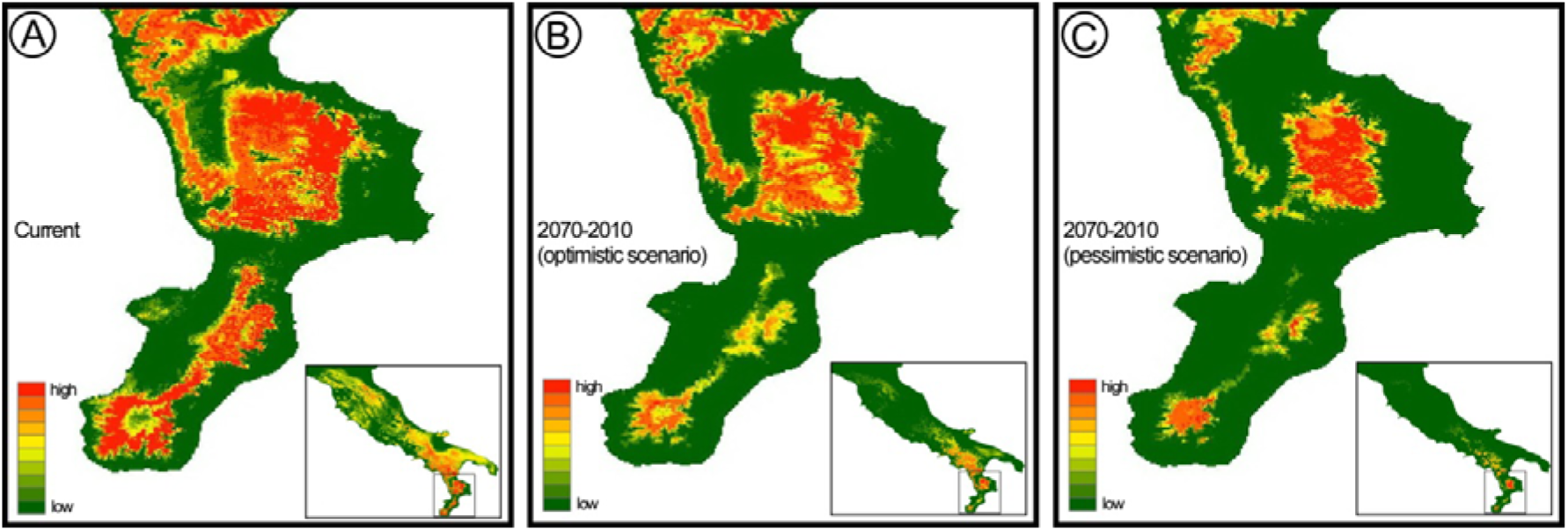
Species distribution model estimated for *B. pachypus* under current bioclimatic conditions (A), and its projection to the near future (2070-2100), under optimistic (B) and pessimistic (C) scenarios of greenhouse gas emission (RCP2.6 and RCP8.5 respectively).

## 4. Discussion

Identifying hotspots of intraspecific genetic diversity is now increasingly seen as a key step towards conservation practices effective in the long-term (Brooks et al., 2015;Thomassen et al., 2011). Our results emphasize that, however, this fundamental step does not suffice, and that finer spatial and temporal scales of analysis, which are characteristic of monitoring programs (Brodersen and Seehausen, 2014; Mimura et al., 2017), should be adopted in order to provide conservation stakeholders with important knowledge to best define conservation priorities and management practices.

The hotspot of intraspecific genetic diversity of *B. pachypus* is not a homogeneous geographic unit, either in terms of genetic diversity, or in terms of species presence and risk factors’ distribution. Moreover, our results provide evidence of geographically structured temporal changes in the analysed features. In the next sections we will discuss these spatio-temporal patterns of variation, and how they could help to identify priorities in the context of both short-term and long-term conservation programs.

### 4.1 Spatial and temporal patterns of variation

Over less than 15 years since the last assessment of species presence (Barbieri et al., 2004), patterns of *B. pachypus* occurrence in southern Italy have dramatically changed, although with marked differences among sub-regions. At the extreme of these differences are the Sila and the Aspromonte massifs. Throughout the Sila region we completely failed to identify sites of current *B. pachypus* presence. While this lack of evidence is not a conclusive argument that the species completely disappeared from the area, it clearly testifies to a dramatic decline. On the other hand, in the Aspromonte region several sites of historical presence were confirmed, and new ones identified. Together with the higher number of individuals per site observed in this area than elsewhere, the Aspromonte region has clearly appeared as the least affected by population declines. Further indication of diffused declines comes also from the observed temporal decreases in genetic diversity estimates, suggesting widespread bottlenecks in population size throughout the hotspot area in southern Italy (Figure 1C).

These findings rise worrisome questions for the conservation of this endangered toad. Why did *B. pachypus* start to decline even in southern Italy? And why is there a geographic structure in this trend? First of all, our results allow us to exclude some of the most commonly invoked answers to these and similar questions. Habitat degradation is probably not a factor in this trend. The Aspromonte and Sila regions are largely covered by national parks, while the Serre region falls within a regional park. Also, in most instances, no evidence of alteration of the physical habitat or of the ecological community was observed at the sites visited. Thus, while a role in species decline at single sites cannot be excluded (Barbieri et al., 2004), habitat degradation can hardly explain the general pattern of declines throughout the study area. Likewise, bioclimatic factors are unlikely to explain the pattern, at least as the single factor. Indeed, the Sila massif, where no site of presence was found, does not suffer of a limited bioclimatic suitability, as compared with neighbouring areas, and most historical sites of presence are located exactly within the area of bioclimatic optimum. Finally, not even *B. dendrobatidis* can be reliably invoked as the ‘lone-killer’ (Pounds and Coloma, 2008 and references therein). As a matter of fact, *B. dendrobatidis* was widespread in the area (including the Aspromonte) at least since the early ‘80s (Canestrelli et al., 2013), that is, well before the observed declines begun. Nonetheless, its role in the observed declines in conjunction with other factors cannot be excluded. Indeed, contrary to results of the previous assessment, *B. dendrobatidis* occurrence was confirmed in the northern but not in the southern portion of the study area.

These results lead to at least two non mutually-excluding hypotheses, useful to attempt answering the above questions. First, the overall reduction of the chytrid geographic distribution [and its absence from sites that are very close to those where high chytrid prevalence was reported until 2003] might be the outcome of epidemic dynamics related to environmental changes. In fact, experimental evidence showed chytrid pathogenicity to decrease at increasing temperatures, with possible implications even at microgeographic scales (Greenspan et al., 2017a), as well as increased host vulnerability to climate change driven by infection (Greenspan et al., 2017b).Second, geographic differences in the mechanisms of resistance of the amphibian host might be implicated (Luquet et al., 2012). In this regard, it is worth recalling that geographic patterns of variation in *B. pachypus* presence and chytrid distribution, are congruent with the geographic structure of the genetic variation among *B. pachypus* populations. Indeed, all the most abundant populations, currently free from chytrid infection, belong to the southern cluster confined to the Aspromonte area. Although not conclusive in disentangling these two scenarios, our results set the stage for future experimental efforts along this road, based on testable research hypotheses.

An interesting corollary to the first scenario comes from bioclimatic suitability predicted for future scenarios in the Aspromonte region. Indeed, *B. pachypus* climate optimum in the Aspromonte massif was predicted to shift upward in elevation in the near future, possibly leading to a selective scenario with two contrasting forces acting on populations, *B. pachypus* being on the horns of a dilemma. Sites of current presence will likely lose their bioclimatic suitability in the near future, but in a direction predicted to favour the toad host over the chytrid pathogenicity (i.e. increasing environmental temperatures). On the other hand, colonizing sites at higher elevation might allow *B. pachypus* to continue matching its bioclimatic optimum, but at the expense of more favourable climatic conditions for its chytrid pathogen. Intriguingly, although speculative for the time being, this scenario might be experimentally evaluated through monitoring environmental variables, epidemiological parameters, and *B. pachypus* performance traits, among populations from the Aspromonte area. Since this is also the only area where the species appeared relatively abundant, such a monitoring and research effort might be one of the last options available to shed light on the mechanisms behind its decline.

### 4.2 Geographic structure of conservation targets

A thorough understanding of mechanisms underlying species’ decline is an essential step towards better informed conservation efforts, but the observed spatial patterns and temporal variations clearly claim for timely actions, aimed at preventing further populations’ disappearance in the short-term. We see four major implications of our results, useful to identify priorities and to optimize resource allocation for such a ‘short-term’ program.

First, and contrary with what previously inferable, not all populations within the hotspot area should be considered as at high priority for species’ conservation programs in situ. Indeed, in most of this area, declines appeared at an advanced stage (if not ultimate). Since an optimal target for in situ actions should be maximizing the chance of species’ persistence, Aspromonte populations appear the ones and last whose demographic, genetic, and epidemiological features, make them suitable for in situ actions, including stringent protection and monitoring of breeding sites.

Second, in those areas (Catena Costiera, Serre) where the species is still present, but where data suggest strong negative demographic trends, pilot experiments of captive-breeding with mixed stocks (individuals of both local and Aspromonte origin) should be implemented. Under the hypothesis of better resistance performance of Aspromonte populations to the chytrid pathogen, such experiments might favour assisted flows of beneficial alleles within local populations (Jones, 2013). If successful under controlled (semi-natural) experimental conditions, these experiments might provide material for subsequent reintroduction programs aimed at species’ recovery.

Third, our results showed a drastic reduction of the geographic occurrence of the chytrid pathogen on its *B. pachypus* host, if compared to what observed in the recent past (Canestrelli et al., 2013), but they do not imply pathogen decline in the area. Thus, monitoring its occurrence *in situ* both in *B.pachypus* and other potential hosts (Simoncelli et al., 2005; Zampiglia et al., 2013), as well as within environmental matrices, will be a mandatory step of in situ actions, as will be also a proper prophylaxis in ex situ programs.

Fourth, but not least, our results showed a possible drastic reduction of habitat suitability expected by 2070-2100, along with an upward range shift. Conservation plans should consider this expected trend, and should plan actions accordingly. In particular, captive-breeding programs should be designed to assess species performance along elevation clines. Useful approaches in this regard might be the assessment of comparative performances (with and without the chytrid pathogen) in experimental climate chambers, or the use of replicated semi-natural settings located along environmental clines. Such approaches might also provide data of extreme value to help identifying priority sites for future restocking or reintroduction programs of this species, both in southern Italy and elsewhere.

## Acknowledgments

We are grateful to Marco Puleo, Antonio Mancuso, and Antonio Barca, for help with sample collection.

## Supplementary material

Supplementary data to this article can be found online at (to be populated).

